# 3D visualization of SARS-CoV-2 infection and receptor distribution in Syrian hamster lung lobes display distinct spatial arrangements

**DOI:** 10.1101/2021.03.24.435771

**Authors:** Ilhan Tomris, Kim M. Bouwman, Youri Adolfs, Danny Noack, Roosmarijn van der Woude, Sander Herfst, Geert-Jan Boons, Bart L. Haagmans, R. Jeroen Pasterkamp, Barry Rockx, Robert P. de Vries

## Abstract

SARS-CoV-2 attaches to angiotensin-converting enzyme 2 (ACE2) to gain entry into cells after which the spike protein is cleaved by the transmembrane serine protease 2 (TMPRRS2) to facilitate viral-host membrane fusion. ACE2 and TMPRRS2 expression profiles have been analyzed at the genomic, transcriptomic, and single-cell RNAseq level, however, biologically relevant protein receptor organization in whole tissues is still poorly understood. To describe the organ-level architecture of receptor expression, related to the ability of ACE2 and TMPRRS2 to mediate infectivity, we performed a volumetric analysis of whole Syrian hamster lung lobes. Lung tissue of infected and control animals were stained using antibodies against ACE2 and TMPRRS2, combined with fluorescent spike protein and SARS-CoV-2 nucleoprotein staining. This was followed by light-sheet microscopy imaging to visualize expression patterns. The data demonstrates that infection is restricted to sites with both ACE2 and TMPRRS2, the latter is expressed in the primary and secondary bronchi whereas ACE2 is predominantly observed in the terminal bronchioles and alveoli. Conversely, infection completely overlaps at these sites where ACE2 and TMPRSS2 co-localize.

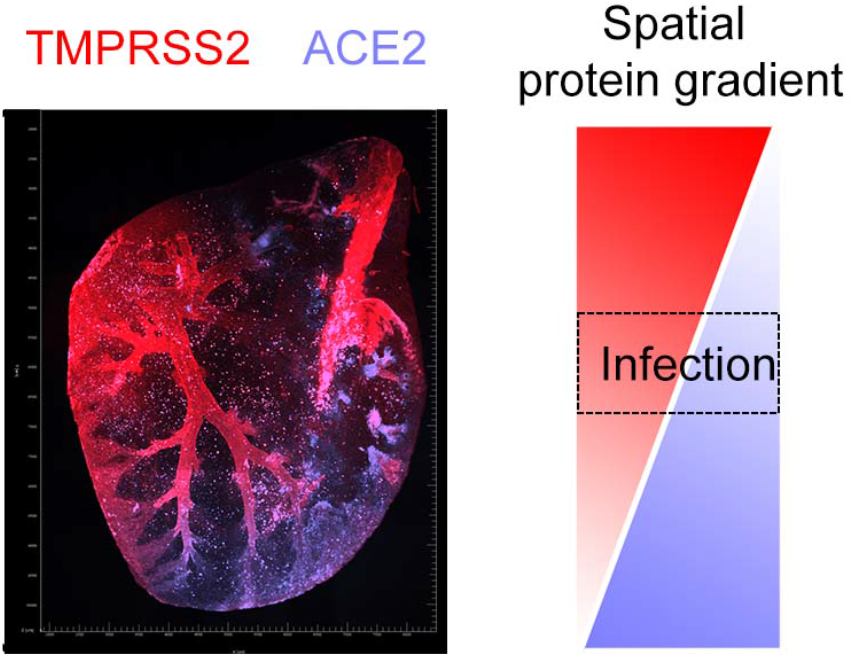

## Main

SARS-CoV-2 has sparked a pandemic and additional means to understand infection dynamics of this virus will facilitate counter-measures. SARS coronaviruses carry a single protruding envelope protein, called spike, that is essential for binding to and subsequent infection of host cells. The trimeric spike protein binds to angiotensin-converting enzyme 2 (ACE2), which functions as an entry receptor for SARS-CoV ^1,2^. After binding and internalization, TMPRSS2 induces the spike protein into its fusogenic form allowing fusion of the viral and target membrane. Several other attachment factors and/or receptors have been reported ^3–6^, but it is generally accepted that ACE2 and TMPRSS2 are essential.

ACE2 and TMPRSS2 are expressed in a wide variety of tissues and have been analyzed using several different genetic techniques ^7–9^. Suprisingly a common denominator in these studies is the high expression of these proteins in extrapulmonary tissues, whereas the virus mainly infects the respiratory tract. A drawback of these genetic studies is that they do not determine biochemical expression and provide limited spatial information. High-resolution mapping of three-dimensional (3D) structures in intact tissues are indispensable in many biological studies, yet hardly employed to study viral receptors in their host organs concerning viral infection. The conventional method of histological sectioning followed by the imaging of individual sections is commonly used and rather valuable, but this process does not provide spatial information. Recent developments in whole organ clearing, imaging, and analysis of these large data sets do now allow for the characterization of whole organs ^10–12^.

Different animal models have been employed to recapitulate SARS-CoV-2 infection in humans, these are instrumental and indispensable to develop vaccines and therapeutics ^13^. Several reviews are available that succinctly compare the advantages and disadvantages of different animal models ^14,15^, and the Syrian hamster is now widely accepted as an extremely suitable small animal model ^16–18^.

Here we aimed to elucidate receptor distribution in the lungs of Syrian hamsters and correlate these patterns with the location of infection. To do so we stained whole lung lobes of SARS-CoV-2 infected Syrian hamsters and control animals, using a variety of antibodies against ACE2, TMPRRS2, and the viral nucleoprotein to detect the accepted functional receptor, restriction factor, and location of infection. The results clearly show that ACE2 and TMPRSS2 are unequally distributed in the lung and that only overlapping regions are infected by SARS-CoV-2.

## Results

### Analysis of SARS-CoV-2 binding and receptors on lung tissue slides of Syrian hamsters lungs

We initially started our studies into receptor binding of SARS-CoV trimeric receptor binding domain (RBD) proteins and the detection of ACE2 in serial tissue slices of Syrian hamster lungs ^19^. We now extended these studies by detecting TMPRSS2 to determine where ACE2 and TMPRRS overlap and if infection would occur in those portions of the lung. SARS-CoV-2 RBDs bind to the apical side of the bronchioles and alveoli of Syrian hamster tissue slides (Fig. 1 A1-3). In Syrian hamster lungs 4-days post-infection (dpi, SARS-CoV-2 infected) SARS-CoV-2 RBD failed to bind as previously shown (Fig. 1 A4-6), whereas significant ACE2 expression is observed in the alveoli and bronchioles of mock and 4-dpi Syrian hamster lungs (Fig. 1 B1-6). Even though we observe alveolar staining of ACE2 in infected Syrian hamster lungs bronchiolar staining is predominant (Fig. 1 B4-6), whilst in non-infected Syrian hamster, the signal is accumulated in both the bronchioles and alveoli (Fig. 1 B1-3). We next characterized TMPRSS2 expression; bronchiolar and alveolar staining was observed in non-infected Syrian hamster lungs, for infected Syrian hamster lungs staining is predominantly visualized in the bronchioles with minor alveolar staining (Fig. 1 C1-6). For infected Syrian hamster lungs an overall reduced signal intensity is observed, in line with previous reports that demonstrate a possible downregulation of receptors and restriction factors ^19,20^. Further staining was performed with an anti-keratin antibody, as keratin 8 (K8) and keratin 18 (K18) are expressed in human alveolar and bronchial epithelial cells ^21,22^. We used this antibody to assess bronchiolar and alveolar staining of TMPRSS2, ACE2, and trimeric SARS-CoV-2 RBDs (Fig. 1 D1-6). In non-infected tissue slides K8/K18 is present in the alveoli and bronchioles (Fig. 1 D1-3), in infected Syrian hamsters a distinct reduction of K8/K18 is observed, probably due to pulmonary damage (Fig. D4-6). Thus, we can detect SARS-CoV receptor binding related to ACE2 and TMPRSS2 specifically in different parts of the bronchiolar and alveolar system. However, the use of tissue slides does not provide spatial information and prompted us to explore volumetric reconstructions instead.

**Fig. 1.**
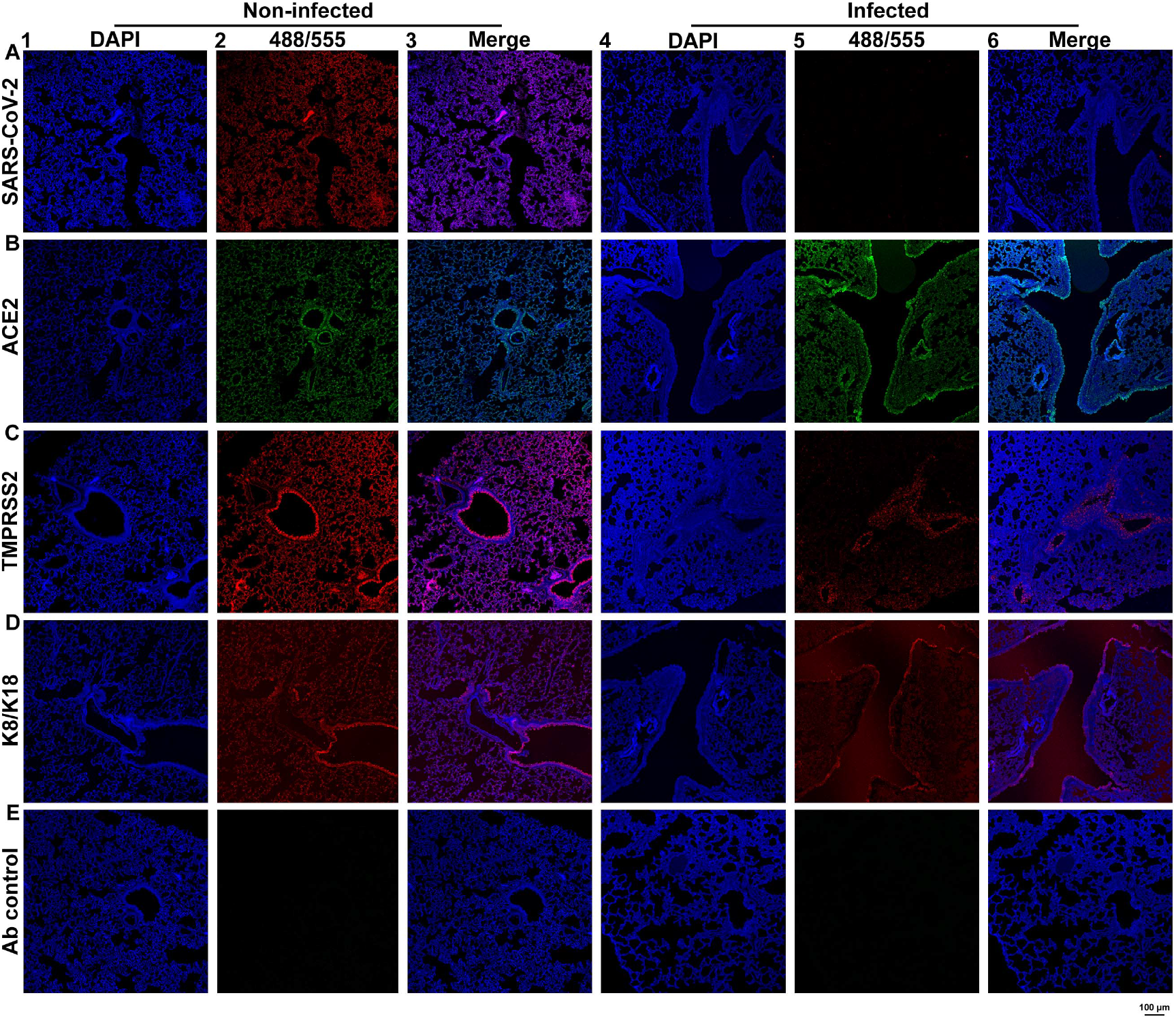
Expression of coronavirus receptors and co-factors on tissue sections of Syrian hamster lung. **(A)** SARS-CoV-2 binding detected on the apical side of the bronchioles and alveoli of non-infected Syrian hamster lungs (1-3), reduced binding on infected Syrian hamster lungs (4-6). **(B)** ACE2 expression observed in Syrian hamster lungs. In infected Syrian hamster lung tissue reduced staining is visible. **(C)** TMPRSS2 expression is characterized, with reduced TMPRSS2 stain in the alveoli of infected Syrian hamsters. **(D)** K8/K18 stains with predominant bronchiolar staining **(E)** With the secondary antibodies only, no staining is observed.

### Syrian hamster lungs abundantly express SARS-CoV-2 receptors and restriction factors in a distinct spatial morphological manner

Volumetric analysis was performed on Syrian hamster lung lobes to assess spatial ACE2 and TMPRSS2 expression in organ samples. The workflow of volumetric imaging and data analysis is described in Figure 2A. To determine SARS-CoV-2 receptor distribution through the Syrian hamster lung, we started with our fluorescent trimeric RBD proteins 19. The trimeric SARS-CoV-2 RBD bound throughout the lung lobes but does not bind the vascular system. We observe distinct signal in the tertiary bronchus, all the way to the alveolar sacs (Fig. 2 B1-4). In the 3D render and orthoslice of trimeric SARS-CoV-2 RBD stained samples (Fig. 2 B1 and B3), we observe a lack of signal in the primary/secondary bronchus relative to the tertiary bronchus. ACE2 receptor expression was highly similar to the SARS-CoV-2 RBD staining (Fig. 2 C1-4), however with varying signal intensity in the secondary, tertiary bronchi, and bronchioles (Fig. 2 C1 and C3). Additionally, comparatively lower antibody staining intensity is observed in the alveoli compared to the SARS-CoV-2 RBD trimer, which is bound to a wider variety of cells. This might be attributed to interactions with other attachment factors, for example, heparin sulfates and sialic acids ^4–6^. Hamster lung lobes incubated with anti-TMPRSS2 antibodies display intense staining in the secondary and tertiary bronchi with minor staining visible in the alveoli and significant staining of the outer regions of the lung (Fig. 2 D1-4). In Fig. 2 E1 and E3 high signal intensity in the primary and secondary bronchus is observed with an anti-K8/K18 antibody, with occasional staining in several tertiary bronchi (Fig. 2 E1-4). In the macroscopic 3D render and orthoslice (Fig. 2 E1 and E3) we identify regions with alveolar staining, this can also be observed with the higher resolution data in Fig. 2 E2 and E4. Due to the extended antibody incubation periods using whole tissues, we used a mix of secondary antibodies only, in which we observe very minor background on the sides of tissues, indicating a lack of a-specific binding while tissue penetration occurs when using primary antibodies (Fig. 2 F1-4). In supplementary Fig 1, the light-sheet data is presented without the autofluorescence channel. Conclusively we observe different spatial expression patterns, which encouraged us to expand our approach by analyzing lung lobes for ACE2 and TMPRRS2 simultaneously.

**Fig. 2.**
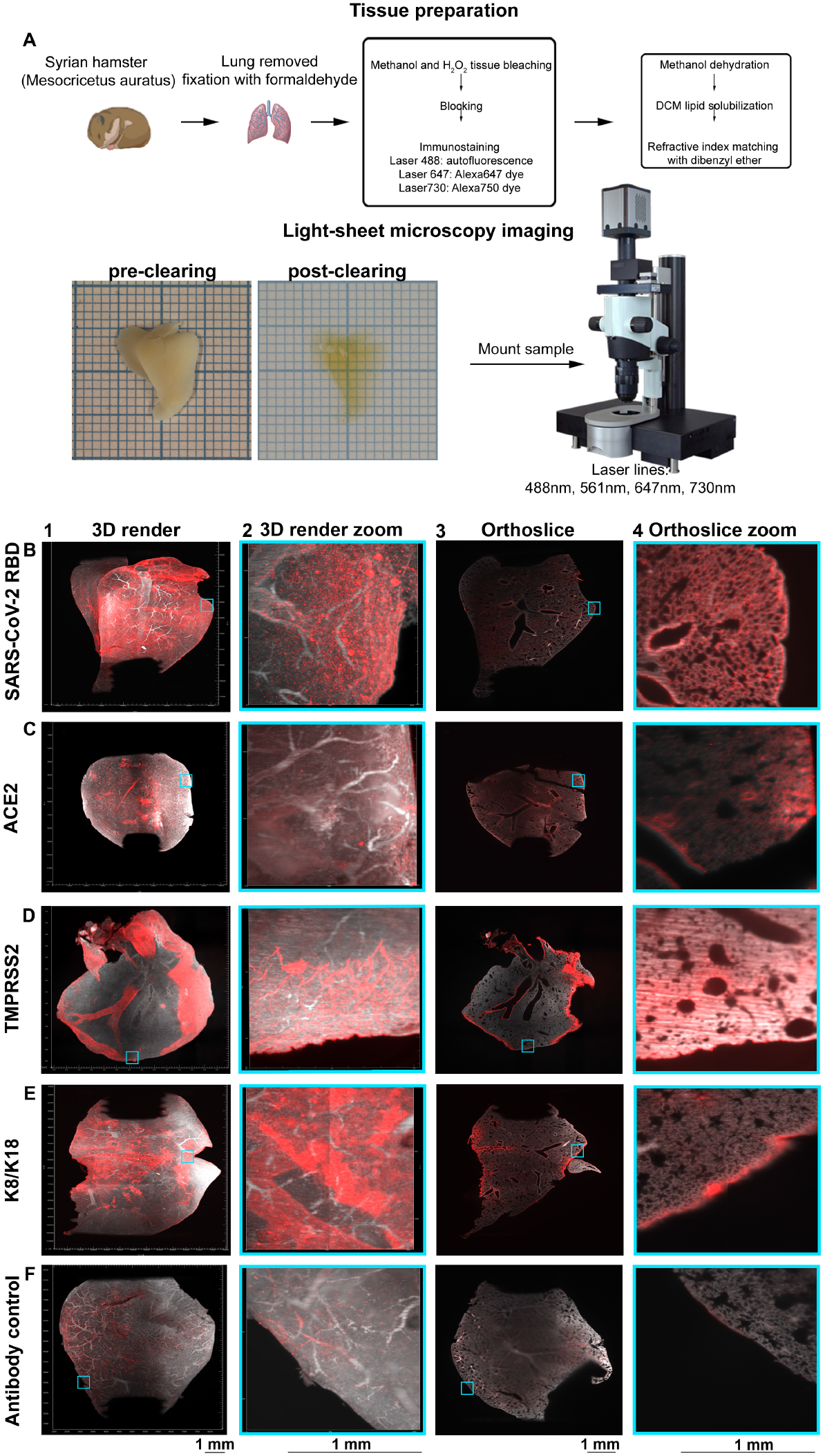
Volumetric analyzes of trimeric SARS-CoV-2 RBD binding and ACE2, TMPRRS2 in non-infected Syrian hamster lung lobes delineate expression patterns. **(A)** Lung lobes from Syrian hamsters are extracted and fixated in 10% formalin. Tissue preparation is performed with bleaching, followed by blocking, immunostaining, dehydration, lipid solubilization and refractive index matching. Refractive index matching for increased light penetration (pre-clearing vs. post-clearing on 1 mm paper), after which imaging is performed and data is analyzed. Orthogonal slices are generated with ImageJ and 3D renders with Imaris. Autofluorescence in grey (488 channel) and staining in red (647 channel). **(B)** Binding of recombinant trimeric SARS-CoV-2 throughout the lung, fluorescence signal is detected in tertiary bronchi and alveoli, without significant signal in the primary/secondary bronchi. **(C)** ACE2 is similarly distributed over the lung as trimeric SARS-CoV-2, with nonuniform signal in the tertiary bronchi, bronchioles and alveoli. **(D)** Significant TMPRSS2 expression in primary, secondary and tertiary bronchi with minor alveolar presence. **(E)** Intense K8/K18 staining in the primary and secondary bronchi, with occasional signal in tertiary bronchi, bronchioles and alveoli. **(F)** Autofluorescence in the 647 channel and minor staining of the outer regions of the lung for the secondary antibody control, pattern does not overlap with previous stains.

### ACE2 and TMPRRS2 are unequally distributed in Syrian hamster lung lobes, displaying a bottom-to-top and top-to-bottom expression profile

To assess where potential viral binding and membrane fusion can occur, co-localization of ACE2 and TMPRSS2 was determined in Syrian hamster lung lobes. Substantial staining with anti-TMPRSS2 antibody in the secondary and several tertiary bronchi is observed (Fig. 3 A1 and supplementary Fig. 2). Signal spreading to the bronchioles and alveoli can also be detected with a similar pattern to previous single stains with minor alveolar staining at the transitioning point from bronchioles to the alveolar ducts (Fig. 3 B1 and supplementary Fig. 2 B). ACE2 expression in this double-stained non-infected Syrian hamster lung appears to be predominantly in tertiary bronchi with minor staining present in the primary/secondary bronchi (Fig. 3 A2 and supplementary Fig. 2 A). During both single and double stainings expression of ACE2 is mainly present in the bronchioles and alveoli (Fig. 3 B2 and supplementary Fig. 2 B), (Fig. 2 B2 and B4). There is a significant overlap of TMPRSS2 and ACE2 in the tertiary bronchi and bronchioles (Fig. 3 A3/4, B3/4, supplementary Fig. 2 A/B, and supplementary video 1/2). In several regions in Fig. 3 A3/A4 and supplementary Fig. 2 A no apparent overlap can be visualized, such as the primary/secondary bronchi where TMPRSS2 expression can be seen, and conversely in several bronchioles/alveolar ducts with ACE2 expression. Thus ACE2 and TMPRSS2 are differentially expressed in the Syrian hamster lung lobes with a top-to-bottom gradient of TMPRSS2 and a bottom-to-top gradient of ACE2.

**Fig. 3.**
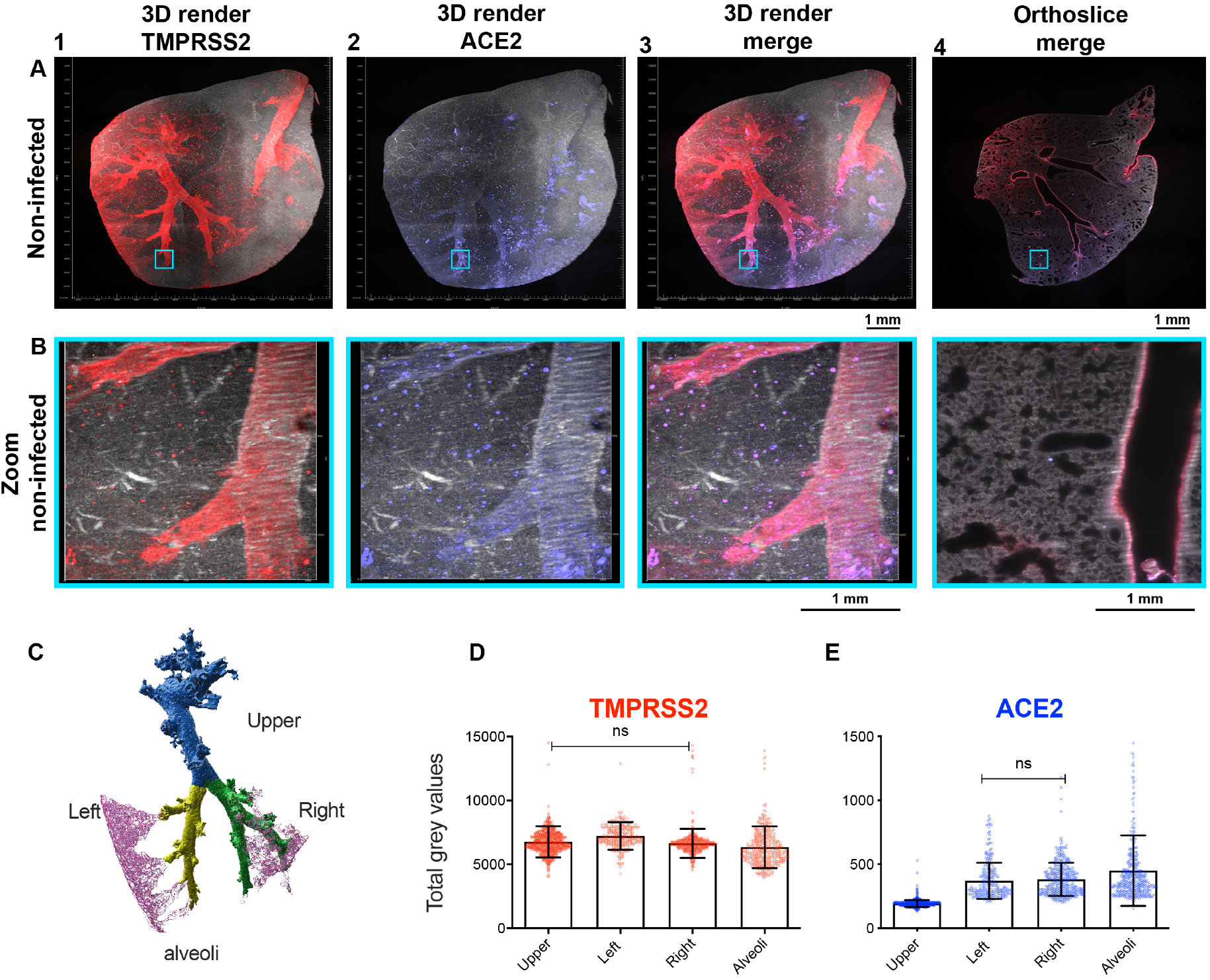
ACE2 and TMPRSS2 partially overlap. Autofluorescence in grey (488 channel), co-staining in red (647 channel) and purple (730 channel), with overlapping regions in pink (colorblind-proof). **(A)** TMPRSS2 expression is seen in the primary/secondary bronchi and several tertiary bronchi, with signal spreading to the bronchioles. ACE2 is predominantly located in the tertiary bronchi with relatively lower signal in the primary and secondary bronchi. A top-to-bottom expression gradient is present for TMPRSS2 and bottom-to-top for ACE2. Overlap in several regions can be observed. 0.63 zoom, voxel resolution X,Y,Z: 4.79 µm, 4.79 µm, 5 µm. **(B)** Minor alveolar TMPRSS2 expression and mostly present at the transitioning point of the bronchioles to the alveolar sacs, a similar pattern is also observed for ACE2 expression. The expression profiles of TMPRSS2 and ACE2 overlap in the bronchioles and alveoli. 6.3 zoom, voxel resolution X,Y,Z: 0.48 µm, 0.48 µm, 2 µm. **(C)** Positions chosen for quantification, 4 different branches and alveoli are indicated. **(D)** Quantification of TMPRSS2 signal in a non-infected Syrian hamster lung lobe from Fig. 3. **(E)** Quantification of ACE2 signal in a non-infected Syrian hamster lung lobe. MP4 files of 3D render and orthoslice merge are provided.

We wondered if our assessment is statistically significant. Indeed, after quantification of TMPRSS2 and ACE2 signal in various regions in the lungs, we observe a top-to-bottom gradient for TMPRSS2 and a bottom-to-top gradient for ACE2 expression (Fig. 3C). TMPRSS2 signal is predominantly present in the larger branches, with decreasing signal intensity in the alveoli (Fig. 3D). ACE2 signal is relatively lower in the larger branches in comparison to the alveoli (Fig. 3E).

### SARS-CoV-2 infected cells in whole Syrian hamster lung lobes spatially resolved

To determine in which specific lung regions SARS-CoV-2 infection takes place, we used hamster lung lobes isolated at four days post-infection, and stained these with anti-NP antibodies. High signal intensity is observed in the bronchi and bronchioles with several tertiary bronchi remaining unstained (Fig. 4 A1 and A2), with minor fluorescence signal in the alveoli (Fig. 4 A3). Additionally, the staining pattern observed with NP staining is slightly different in comparison to trimeric SARS-CoV-2 RBD staining (Fig. 2 A2 and A4), with trimeric RBD staining being present all over the alveoli whilst anti-NP antibody binding is non-continuous (Fig. 4 A1-3). We also analyzed the anti-NP antibody on non-infected hamster lungs and observed some background at the major bronchi on the outside regions of the lungs, albeit with an apparent lower fluorescence signal relative to the infected Syrian hamster lung (Fig. 4 B1-3 and supplementary Fig. 3 B, 4 A). This was however not the case for other NP antibodies tested, for example, the 40143-R001 and MA1-7401 gave significant background staining through the bronchiolar system (supplementary Fig. 4 B and C). Thus an adequate analysis of commercial antibodies cannot be overstated, as we have seen differential results with anti-ACE2 antibodies previously ^19^. The antibody control (secondary antibodies) for the infected hamster lung samples displays minor background staining without any similarity to the NP stain (Fig. 3 C1-3). Conclusively, we can confidently detect infected regions within the whole lung lobes of infected Syrian hamsters.

**Fig. 4.**
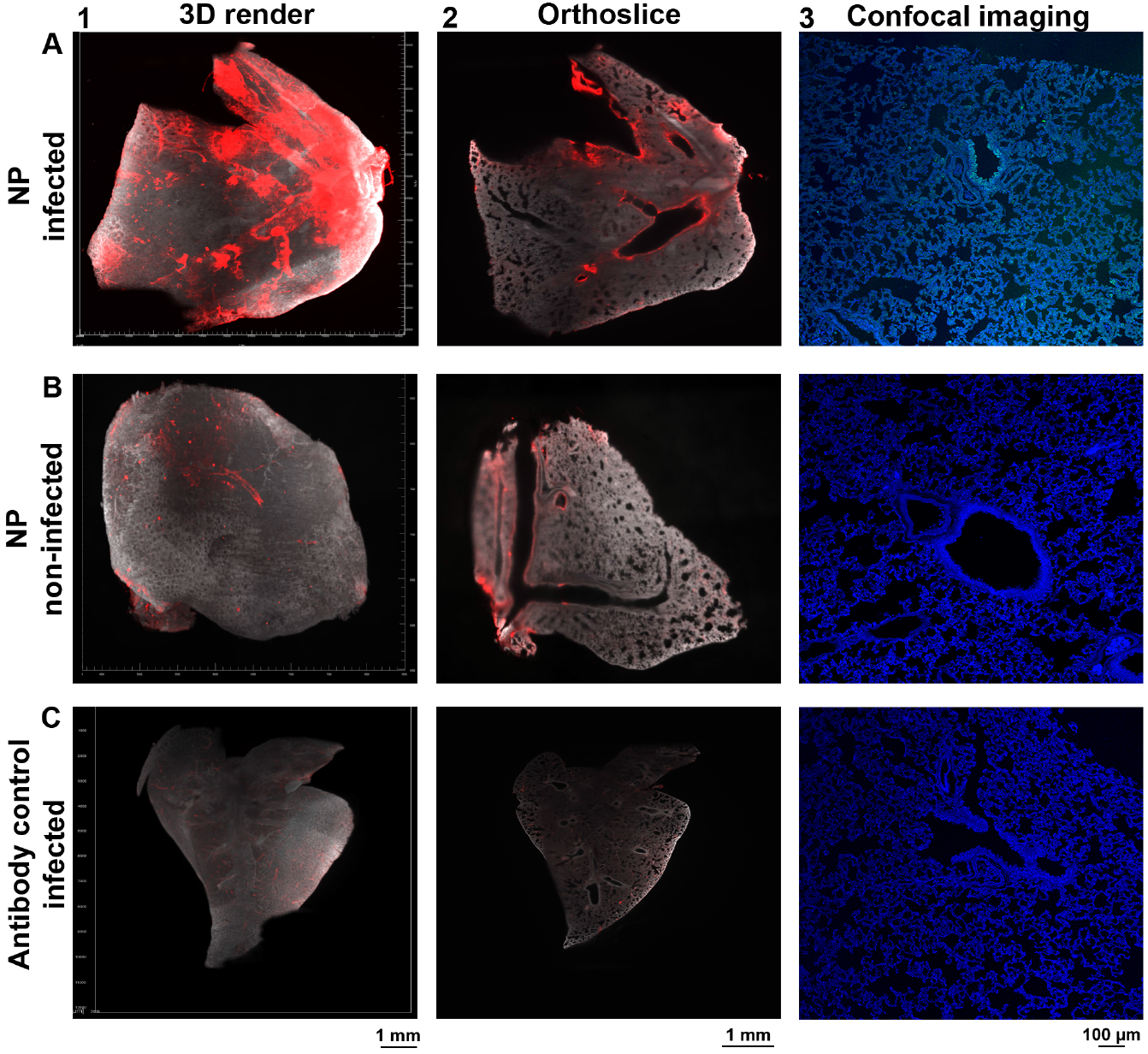
Detecting infected cells within a whole Syrian hamster lung lobe. Autofluorescence in grey (488 channel) and staining in red (647 channel). **(A)** Significant NP staining in the primary/secondary bronchi and bronchioles, with occasional tertiary bronchi and alveolar staining (light-sheet and confocal imaging data). **(B)** A-specific binding in non-infected when using anti-NP antibody in the bronchi and outer regions of the lung, although with a lower apparent signal intensity relative to infected lungs (light-sheet and confocal imaging data). **(C)** Minor autofluorescence and a-specific staining with secondary antibodies, observed pattern does not overlap with previous stains (light-sheet and confocal imaging data).

### Infected Syrian hamster lungs display infection patterns that are retained to locations where both ACE2 and TMPRRS2 co-localize

Now with the ability to detect ACE2, TMPRSS2, and infected cells in a 3D format, we determined the overlap of ACE2 and SARS-CoV-2 infection to assess if infection was completely restricted to regions with ACE2 expression (Fig. 5 A and B). We observed NP staining in several regions of the secondary bronchi extending to tertiary bronchi and bronchioles (Fig. 5 A1 and supplementary Fig. 5 A). The primary bronchi and the other secondary bronchi in the upper portion of the lung lobe appear to be completely clear of infection. NP staining is observed in the alveoli, albeit at a relatively low signal intensity (Fig. 5 B1 and supplementary Fig. 5 B), additionally over the entire lung a large number of foci are present (Fig. 5 B2 and supplementary Fig. 5 B). ACE2 expression is detected again mainly in the tertiary bronchi in the lower parts of the lung lobe (Fig. 5 A2 and supplementary Fig. 5 A). No significant signal is detected in the primary bronchi and the other secondary bronchi also appear to express low levels of ACE2. ACE2 patterns were strikingly similar to the NP patterns, including a major overlap (Fig. 5 A3/B3, supplementary Fig. 5 A/B, and supplementary video 3/4). NP staining appears to overlap significantly with ACE2 staining in the tertiary bronchi, bronchioles, and alveoli (foci). However, ACE2 staining does not completely overlap with NP staining, in the lower regions of infected Syrian hamster lung lobes there appears to be a lack of NP staining whilst significant ACE2 staining is observed. Thus, SARS-CoV-2 infection is restricted to the middle portion of the lung lobe, with receptors in the lower parts not utilized, perhaps due to lower levels of TMPRSS2.

**Fig. 5.**
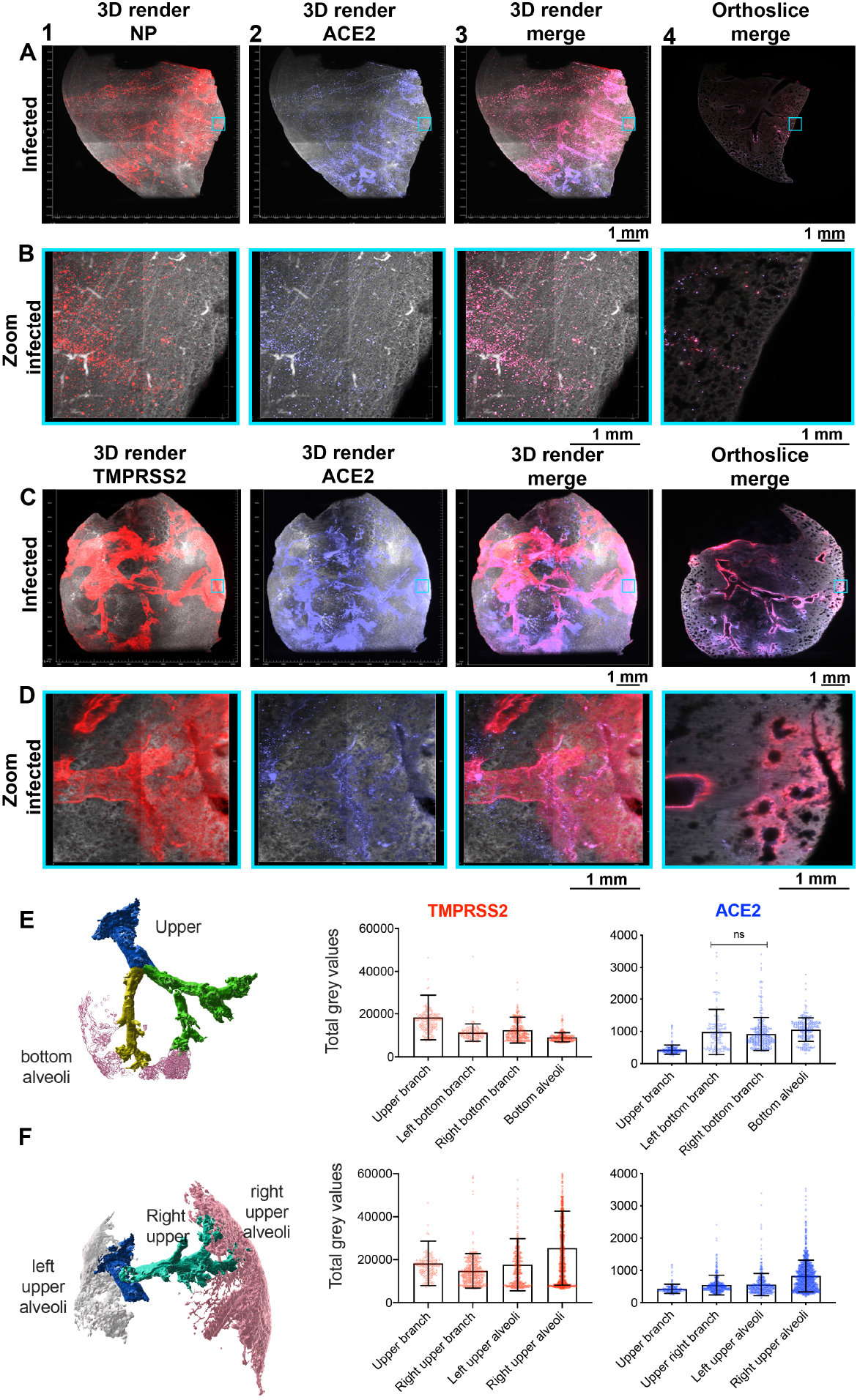
Visualizing tropism of SARS-CoV-2 infection with ACE2 and TMPRSS2 expression patterns at 4-dpi. Autofluorescence in grey (488 channel), co-staining in red (647 channel) and purple (730 channel), with overlapping regions in pink (colorblind-proof). **(A)** In several regions of the lung, NP staining is observed from the secondary bronchi extending to the tertiary bronchi, a top-to-bottom gradient. ACE2 is again a bottom-to-top gradient, with signal in the alveoli and various tertiary bronchi. There is major overlap of NP with ACE2 in the tertiary bronchi, bronchioles and alveoli (foci). In the lower portion of the lung there is no apparent overlap of NP and ACE2. 0.63 zoom, voxel resolution X,Y,Z: 4.79 µm, 4.79 µm, 5 µm. **(B)** In the alveoli there is significant overlap of NP and ACE2, the previously observed foci appear to be alveolar. Several NP stained foci do not superimpose with ACE2 stained foci. 6.3 zoom, voxel resolution X,Y,Z: 0.48 µm, 0.48 µm, 2 µm. **(C)** The expression profile of TMPRSS2 in infected Syrian hamster lungs is highly similar to expression in non-infected lungs. Fluorescence signal is spreading towards the bronchioles. Likewise, ACE2 expression is also highly similar to mock lungs, with protein staining in tertiary bronchi and bronchioles, 0.63 zoom, voxel resolution X,Y,Z: 4.79 µm, 4.79 µm, 5 µm. **(D)** At the transitioning point of bronchioles and alveoli TMPRSS2 and ACE2 superimpose, whilst in the alveoli only ACE2 expression is present. 6.3 zoom, voxel resolution X,Y,Z: 0.48 µm, 0.48 µm, 2 µm. **(E)** Positions chosen for quantification in the infected Syrian hamster lung lobe from panel C, different branches and alveoli are indicated to analyze expression in the bottom part of the lung lobe and the quantification of TMPRSS2 and ACE2 signal in the bottom regions of the infected Syrian hamster lung lobe. **(F)** Positions chosen for quantification in an infected Syrian hamster lung lobe from panel C, different branches and alveoli are indicated to analyze expression in the upper part of the lung lobe and the quantification of TMPRSS2 and ACE2 signal in the upper regions of the infected Syrian hamster lung lobe. MP4 files of 3D render and orthoslice merges are provided.

Next, we analyzed the spatial distribution of TMPRSS2 concerning ACE2 in infected Syrian hamster lungs (Fig. 5 C and D). A similar expression profile is observed compared to the duplicate experiment in non-infected lungs (Fig. 3). Significant TMPRSS2 expression is detected in the primary, secondary and tertiary bronchi, with a spread to the bronchioles and alveoli, similarly to previous single and double stains with alveolar staining at the transitioning point of the bronchioles and alveoli (Fig. 5 C1/D1 and supplementary Fig. 5 C/D). ACE2 expression peaks immensely in the tertiary bronchi, bronchioles, and alveoli (Fig. 5 C2/D2 and supplementary Fig. 5 C/ D), with clearly lower signal in the primary and secondary bronchi (Fig. 5 C2 and supplementary Fig. 5 C). The overlap of ACE2 and TMPRSS2 is predominantly present in the tertiary bronchi and transitioning point of the bronchioles and alveoli (Fig. 5 C3/D3, supplementary Fig. 5 C/D, and supplementary video 5/6). This overlap corresponds with where we observe NP staining, confirming the necessity of both ACE2 and TMPRSS2 for infection.

TMPRSS2 and ACE2 expression gradients in infected Syrian hamster lungs appeared to be near identical to non-infected hamster lungs. Whereas TMPRSS2 expression is predominantly restricted to the bronchi and upper regions of the lung, ACE2 appears to be prominently present lower in the lung lobe. We observed these gradients in multiple different lung lobes with different staining approaches. As for the non-infected lung lobe, after quantification, we observe a similar pattern. We observe a higher TMPRSS2 signal in the upper branches and a significantly lower signal when reaching the “Bottom alveoli” (Fig. 3 C/D/E). This is also the case for ACE2 but then inverted, with decreased ACE2 signal in the “Upper branch” and increased signal towards the “Bottom alveoli”. We also quantified the alveolar signal in other portions of this infected and co-stained lung the alveolar TMPRSS2 signal is dependent on the location and that alveoli in higher regions of the lung lobe may have a higher apparent fluorescence signal (Fig. 5 E and F). In this example the pattern for ACE2 is similar to the pattern observed before, an elevated signal in the alveoli and reduced fluorescence signal in the larger branches (Fig. 3 C/D/E and 5 E/F). We additionally quantified several other staining experiments, ACE2 fluorescence signal in the alveoli of infected hamster lung appears to be similar in terms of fluorescence, with decreasing intensity towards the upper branches (Supplementary Fig. 6 A and B). TMPRSS2 in infected hamster lobes provides a comparable impression by displaying immense fluorescence signal in the “Upper branch” and noticeably decreased signal in the “Bottom-lower branch” (Supplementary Fig. 6 C and D). We, therefore, hypothesize that infection occurs where ACE2 and TMPRSS2 are both present in sufficient density.

## Discussion

In this study, we demonstrate distinct spatial distribution, or gradient, of ACE2 and TMPRSS2 proteins in the lung lobes of Syrian hamsters, which is considered to be the relevant small animal model for studying SARS-CoV-2 infection ^17^. Recently, using light-sheet microscopy, virus infection in the upper respiratory tract of ferrets was shown by detecting NP positive cells, which formed relatively rare foci ^23^. Yet the tissues used in this preprint were significantly smaller compared to the whole lung lobes used here. Importantly we now show the proteinaceous presence of the functional receptor ACE2 ^2^, and the essential restriction factor TMPRSS2 ^24–26^, in concert with NP detection. Importantly, we observed that in lung regions in which ACE2 and TMPRSS2 overlap, actual infection took place.

Ferrets and Syrian hamsters display dramatically different SARS-CoV-2 infection profiles. Where in ferrets infection seems to be restricted to the upper respiratory tract ^27–29^, in Syrian hamsters infection is observed throughout the respiratory tract ^17,30^. In most studies whole Syrian hamster lungs are analyzed to determine where infection occurs, however, even whole lungs were then subjected to cutting to generate tissue slices. Furthermore, virus titers are normally determined from whole organs sections, and thus also fail to give a more detailed overview of infection dynamics. Using whole Syrian lung lobes, we here provide a blueprint to analyze infection patterns, and receptor expression organization in a whole organ section. For SARS-CoV-2 it would be of interest to investigate extrapulmonary infection patterns but we also envision that these approaches can be utilized for other pathogens ^10,31,32^.

An intriguing observation is that in previous normal lung cohorts hardly any ACE2 protein expression was observed in the human lung and bronchus ^33^. Indeed it is known that ACE2 primarily resides in the alveoli, and thus in line with our light-sheet microscopy observations. On the other hand, TMPRSS2 protein expression distribution is hardly known. Transcriptomic data suggests at least low expression levels of ACE2 in respiratory cells and indicates that the co-factor TMPRSS2 is highly expressed with broader distribution ^8^. We demonstrate that the TMPRSS2 co-factor indeed appears to be present with a relatively higher abundance compared to ACE2 (Fig. 4 and 6), albeit being predominantly restricted to the major lung branches in Syrian hamsters. The fact that several studies using anti-ACE2 antibodies appeared not to correlate with infection patterns ^33,34^, and our observation of broad SARS-CoV-2 trimeric RBD binding, indicates that infection is a process with multiple complexities.

Finally, it might be possible to extend our data into the increased ACE2 receptor binding affinities of circulating variant viruses ^35–37^. A hypothesis would be that these viruses can infect the upper respiratory tract, in which ACE2 is scarce, more efficiently leading to increased transmission, a highly similar model as observed for human influenza A viruses ^38,39^. The protein gradient of ACE2 and TMPRSS2 can also be discussed to this extend, whereas for a2-6 linked sialic acids, the receptor for human influenza viruses, also displays a top to bottom gradient^40^. However, the biological function of these gradients remains to be elucidated.

## Methods

### Animal tissues and ethical statement

Animals were handled in a BSL3 biocontainment laboratory. Animals were housed in groups of 2 animals in filter top cages (T3, Techniplast), in Class III isolators allowing social interactions, under controlled conditions of humidity, temperature, and light (12-hour light/12-hour dark cycles). Food and water were available ad libitum. Animals were cared for and monitored (pre-and post-infection) by qualified personnel. The animals were sedated/anesthetized for all invasive procedures.

### Animal procedures SARS-CoV-2

Female Syrian golden hamsters (Mesocricetus auratus; 6-week-old hamsters from Janvier, France) were anesthetized by chamber induction (5 liters 100% O_2_/min and 3 to 5% isoflurane). Animals were inoculated with 10^5^ TCID50 of SARS-CoV-2 or PBS (mock controls) in a 100 μl volume via the intranasal route. Animals were monitored for general health status and behavior daily and were weighed regularly for the duration of the study (up to 22 days post-inoculation; d.p.i.). Animals were euthanized on day 4 after inoculation, and lung samples were removed and stored in 10% formalin for histopathology.

### Antibodies

ACE2 antibody abcam, ab27269 1:400

TMPRSS2 antibody Santa Cruz sc-515727 1:400

Anti-NP, sino-biological 40143-MM05 1:400

Anti-NP, sino-biological 40143-R001 1:400

Anti-NP, thermofisher MA1-7401 1:400

K8/K18, progen, 90001 1:400

Donkey-anti-rabbit750, abcam ab175731 1:500

Donkey-anti-GP647, jackson immunoresearch, 706-605-148 1:800

Goat anti-rabbit647, thermofisher A-21245 1:800

Goat-anti-mouse647, thermofisher, A-21235 1:800

Strepmab-HRP, iba-lifesciences 2-1509-001

### Tissue staining

Sections of formalin-fixed, paraffin-embedded Syrian hamster lungs were obtained from the department of Viroscience, Erasmus University Medical Center, The Netherlands. Tissue sections were rehydrated in a series of alcohol from 100%, 96% to 70%, and lastly in distilled water. Tissue slides were boiled in citrate buffer pH 6.0 for 10 min at 900 kW in a microwave for antigen retrieval and washed in PBS-T three times. Endogenous peroxidase activity was blocked with 1% hydrogen peroxide for 30 min. Tissues were subsequently incubated with 3% BSA in PBS-T overnight at 4 °C. The next day, the purified viral spike proteins (50 μg/ml) were added to the tissues for 1 h at RT. With rigorous washing steps in between the proteins were detected with mouse anti-strep-tag-antibodies (IBA) and goat anti-mouse IgG antibodies (Life Biosciences).

### IDISCO

SARS-CoV-2 infected and non-infected Syrian hamster lungs were provided in 10% formalin, for longer storage hamster lungs were kept in PBS and 0.01% sodium azide. Dehydration of the lungs was performed by washing with PBS for 1.5 hours, followed by 50% methanol for 1.5 hours, 80% methanol for 1.5 hours, and as of last 100% methanol for 1.5 hours on a tilting laboratory shaker. Bleaching was performed overnight at 4 °C in 90% methanol (100% v/v) and 10% H_2_O_2_ (30% v/v). Tissues were rehydrated with 100% methanol for 1 hour, followed by 100% methanol for 1 hour, 80% methanol, 50% methanol, and 1x PBS for 1 hour on a tilting shaker. Syrian hamster lungs were subsequently blocked for 24 hours at room temperature (22 °C) in 1x PBS with 0.2% gelatin, 0.5% triton-x-100, and 0.01% sodium azide (PBSGT) on a tilting shaker. Hamster lungs were stained with trimeric SARS-CoV-2 RBD protein and/or with primary/secondary antibodies possessing an alexa647 or alexa750 dye. After blocking samples were incubated for 120 hours with primary antibody or trimeric RBD protein in PBSGT with 0.1% saponin (S2149, Sigma-Aldrich) on a shaking incubator at 200 rpm, following 120 hours incubation, hamster lungs were washed with PBSGT six times for 1 hour on a tilting shaker. Post-PBSGT wash, samples stained with primary antibodies were incubated with secondary antibodies that possess alexa647 or alexa750 dyes, the antibodies were diluted in PBSGT with 0.1% saponin and filtered with 0.45 µm filter for potential antibody aggregates. Hereafter samples were incubated for 120 hours at room temperature on a shaking incubator at 200 rpm. Samples stained with trimeric SARS-CoV-2 RBD were incubated with primary antibody strepmabHRP (2-1509-001, IBA lifesciences) with specificity towards the TwinStrep-tag at the C-terminus of the trimeric protein for 120 hours in PBSGT with 0.1% saponin. Following secondary antibody staining and tissues treated with trimeric RBD proteins, the hamster lungs were washed with PBSGT six times for 1 hour on a tilting shaker. Secondary antibody staining was also performed for trimeric RBD stained sample either using antibodies with alexa647 or alexa750 dyes with an incubation period of 120 hours, after staining washing was performed with PBSGT six times for 1 hour on a tilting shaker. Immunostained samples were subsequently treated with 50% methanol for 24 hours, 80% methanol for 24 hours, 100% methanol for 24 hours followed by 100% methanol for 24 hours on a tilting shaker for dehydration of the lungs. Hereafter lipid solubilization was performed with dichloromethane (Thermofisher, 402152) for 40 minutes after which refractive index matching and optical clearing were performed overnight with dibenzyl ether (108014, Sigma-Aldrich).

### Light-sheet microscopy

Light-sheet imaging was performed using the Ultramicroscope II (LaVision BioTec) equipped with an MVX-10 Zoom Body. The laser lines 488 nm (Coherent OBIS 488-50 LX Laser 50mW), 647 nm (Coherent OBIS 647-120 LX Laser 120mW), and 730 nm (Coherent OBIS 730-30 LX Laser 30mW) were used with 70% laser power for 488 channel, 70% laser power for the 647 channel and 100% laser power for the 730 nm channel. Emission filters ET525/50 (488 channel), ET676/29 (647 channel) and 716/40 (730 channel) filter was used with a Neo 5.5 sCMOS detector (2560×2160 pixels, pixel size: 6.5 × 6.5 µm^2^). A corrected dipping cap (CDC) (1.26x – 12.6x) was utilized with an Olympus MVPLAPO 2x objective lens. Imaging was performed with 0.63 zoom and step size of 5 µm (voxel resolution X, Y, Z: 4.79 µm, 4.79 µm, 5 µm), also with 6.3 zoom and step size of 2 µm (voxel resolution X, Y, Z: 0.48 µm, 0.48 µm, 2 µm). The exposure time was set to 200 ms and illumination for the 0.63 zoom was bidirectional and 6.3 zoom unidirectional. The sheet numerical aperture was 0.033 with a sheet thickness of 7.09 µm and sheet width of 40%. The imaging chamber was filled with dibenzyl ether and the microscope software is Inspector version 7.1. Obtained image stacks were analyzed with ImageJ v1.54f and Imaris 9.6. Stills of image slices (orthoslices) were generated in ImageJ, whilst 3D renders were generated in Imaris. TIFF files generated by the light-sheet microscope were imported to ImageJ with “File -> Import -> Bio-formats”, stack is viewed as “Hyperstack” color mode was set to “Composite” and “Display metadata” with “Display OME-XML metadata” was enabled to obtain voxel information. Brightness adjustments were performed for each channel with setting “Image -> Adjust -> Brightness/Contrast”. The 488 channel color was set to hex code #FFFFFF, the 647 channel was set to hex code #FF0000 and the 730 channel was set to hex code #8080FF. Snapshots of the Z-stack were saved as JPEGs and the orthoslice animations with “Save As -> Avi -> Compression JPEG and Frame Rate 30 FPS”. For the 3D renders obtained imaging data was imported into Imaris, display adjustments were made for each channel. Snapshots and animations were made with 3D renders (3D View) within Imaris. Volume mode was set to maximum intensity projection (MIP) and the rendering quality was set to highest. Frame settings were used for the outer bounding box of the 3D render, “Box”, “Tickmarks”, and “Axis labels” were enabled. The spacing of the tickmarks was set to µm 500 for the X, Y, and Z-axis. The 488 channel color was set to hex code #FFFFFF, the 647 channel was set to hex code #FF0000 and the 730 channel was set to hex code #8080FF. The 3D animations were made with 1920 × 1080 resolution (16:9) and 360 frames, 360° horizontal turn.

### Quantification

For TMPRSS2 and ACE2 quantification surfaces and masks were generated in Imaris with the “Surfaces Creation Wizard” (according to the reference manual). In this surface/mask regions of interest were selected, i.e. the branches and alveoli in the hamster lung. Regions of interest were selected in the surface tool with the corresponding source channel, whereby the 647 channel with TMPRSS2 signal was used for TMPRSS2 and ACE2 double-stained samples and 730 source channel for ACE2 and anti-NP double-stained samples. For the TMPRSS2 single stained sample, the 647 channel was used as the source channel. Thresholding (Absolute Intensity) was performed arbitrarily to obtain complete coverage of the alveoli and branches. The automatically provided value for “Sphere Diameter” was used. For the separation of two or more objects that are identified as one, the “Split touching Objects (Region Growing)” setting was utilized. The “Seed Points Diameter” value was set to 35 µm. The generated surfaces were subsequently used to mask the TMPRSS2 and ACE2 channels, after which statistical values were obtained and exported to GraphPad Prism 9.

## Acknowledgments

The authors thank the MIND facility of the UMC Utrecht Brain Center for support with iDISCO.

## Author contributions

I.T., and R.P.dV. designed the project; I.T., K.M.B., Y.A., D.N., R.vd.W., S.H., and B.R. performed experiments and data analysis; G.-J.B., S.H. B.L.H., J.P., B.R., and R.P.dV. provided scientific guidance on the experimental setup and data interpretation; I.T., and R.P.dV. wrote the manuscript, and all authors provided comments and suggestions on the manuscript.

## Data and materials availability

All data is available in the main text or the supplementary materials. Sharing of materials described in this work will be subject to standard material transfer agreements.

## Ethics statement

Research was conducted in compliance with the Dutch legislation for the protection of animals used for scientific purposes (2014, implementing EU Directive 2010/63) and other relevant regulations. The licensed establishment where this research was conducted (Erasmus MC) has an approved OLAW Assurance # A5051-01. Research was conducted under a project license from the Dutch competent authority and the study protocol (#17-4312) was approved by the institutional Animal Welfare Body.

