## Supplemental figures for "3D visualization of SARS-CoV-2 infection and receptor distribution in Syrian hamster lung lobes display distinct spatial arrangements"

#### Supplementary Fig 1

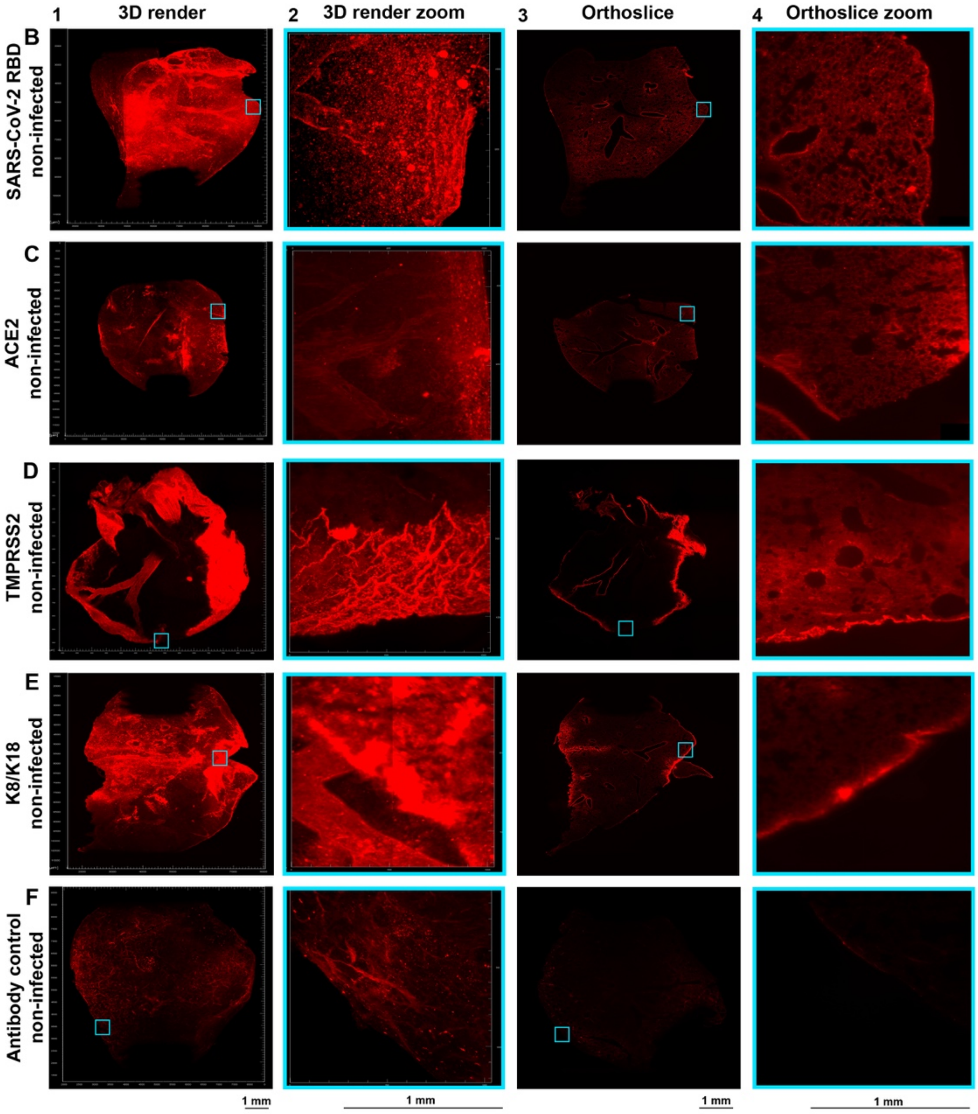


**Supplementary Fig 1. 647 channel only supplementary data of non-infected single stains, related to figure 2.** staining in red (647 channel).

**A)** Recombinant trimeric SARS-CoV-2 binding throughout the lung lobe.

**B)** ACE2 distribution over Syrian hamster lung lobe.

**C)** TMPRSS2 signal present in larger branches.

**D)** K8/K18 staining in the primary, secondar and in tertiary bronchi, with bronchiolar and alveolar staining bronchioles and alveoli.

**E)** Autofluorescence in the antibody control.

25 **Supplementary Fig 2**

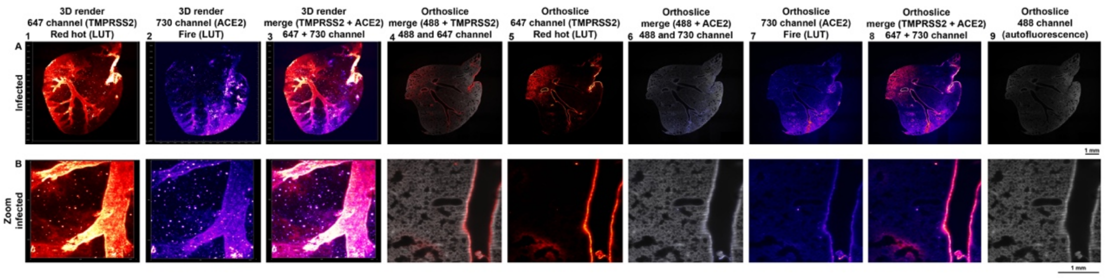


**Supplementary Fig 2. Supplementary analyses of ACE2 and TMPRSS2 in a non-infected Syrian hamster lung, related to figure 3.**

**A)** TMPRSS2 expression in primary, secondary and tertiarty bronchi. Signal spreading to bronchioles. ACE2 staining in tertiary bronchi and bronchioles. In the tertiary bronchi overlap visualised of the receptor and co-factor 0.63 zoom, voxel resolution X,Y,Z: 4.79 µm, 4.79 µm, 5 µm**.**

**B)** Bronchiolar and alveolar TMPRSS2 and ACE2 expression visualized and superimposed. 6.3 zoom, voxel resolution X,Y,Z: 0.48 µm, 0.48 µm, 2 µm.

**Row 1)** 3D render 647 channel (Red hot false color).

**Row 2)** 3D render 730 channel (Fire false color).

**Row 3)** merge 3D render 647 channel (Red hot false color) and 730 channel (Fire false color).

**Row 4)** orthoslice merge 488 and 647 channel.

**Row 5)** Orthoslice 647 channel (Red hot false color).

**Row 6)** orthoslice merge 488 and 730 channel. Row 7) orthoslice 730 channel (Fire false color).

**Row 8)** merge orthoslice 647 channel (Red hot false color) and 730 channel (Fire false color).

**Row 9)** orthoslice 488 channel (autofluorescence)

#### Supplementary Fig 3

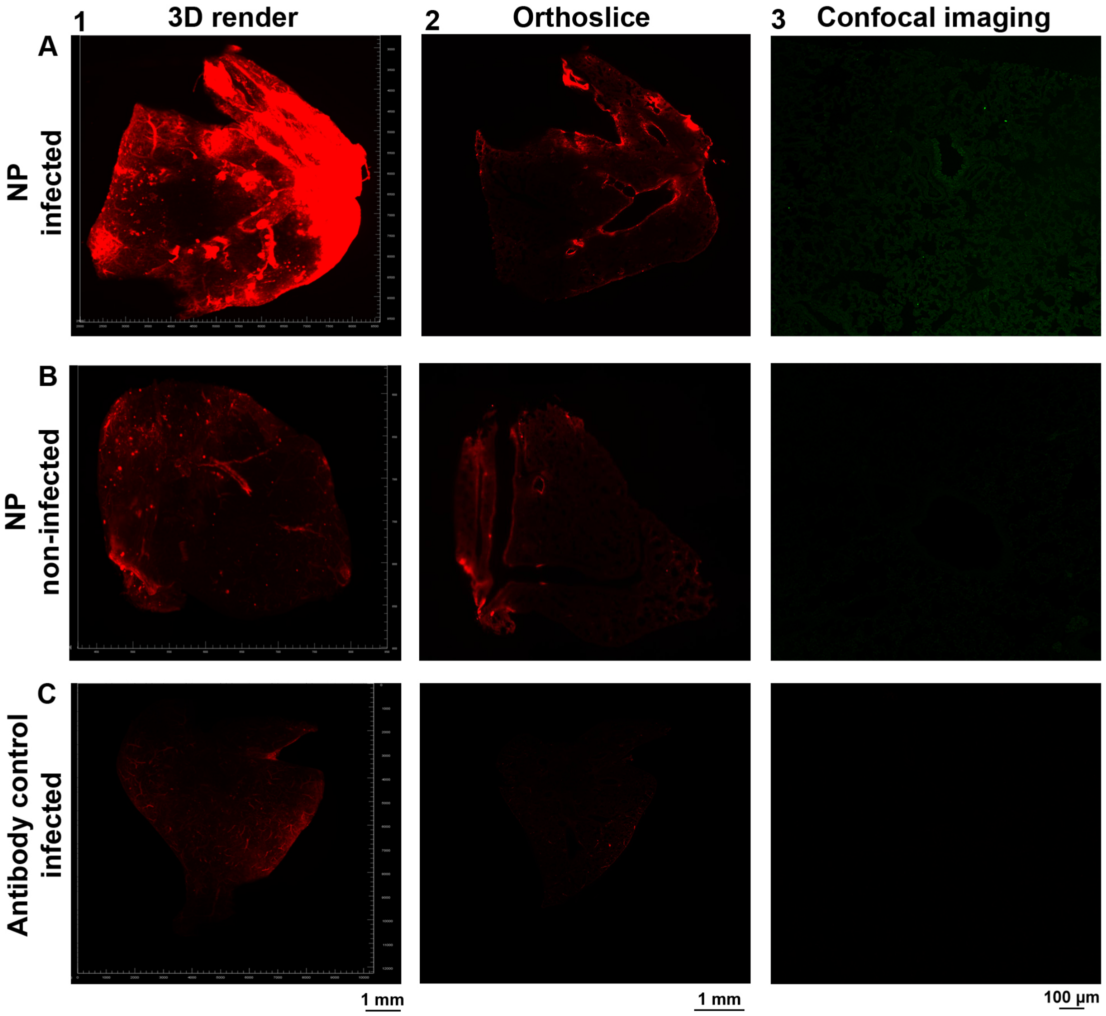


30

**Supplementary Fig 3. 488 and 647 channel only supplementary data of non-infected single stains, related to figure 4.**

Light-sheet staining in red (647 channel) and confocal staining in green (488 channel).

**A)** NP staining in the primary/secondary bronchi.

**B)** Anti-NP antibody displays hardly an aspecific binding in Syrian hamster lung lobes.

**C)** Autofluorescence in the antibody control.

#### Supplementary Fig 4

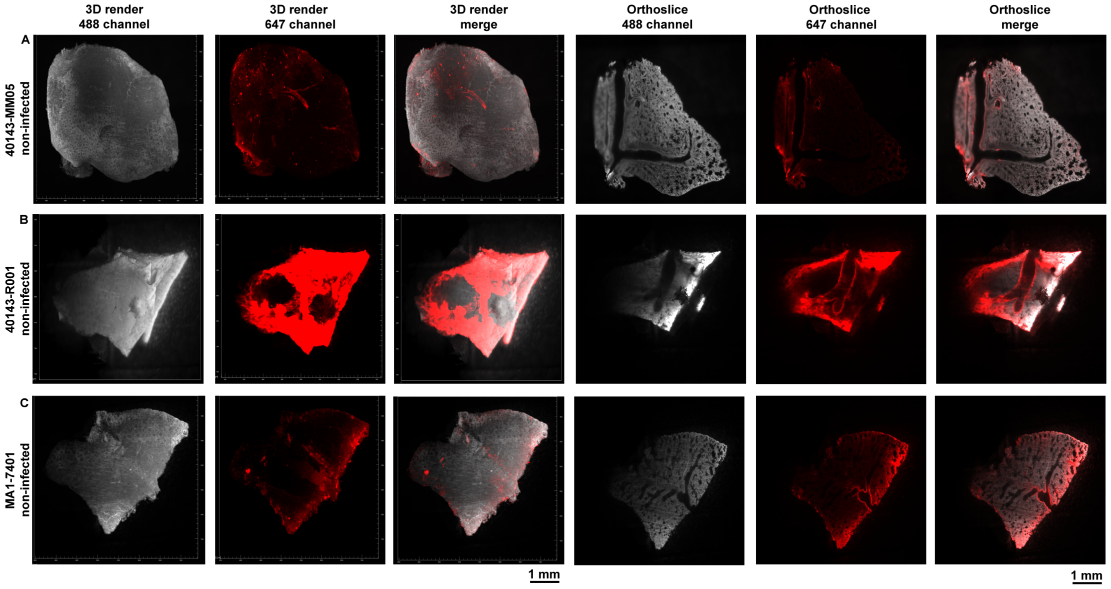


**Supplementary Fig 4. Supplementary analysis of NP antibodies, related to figure 4.**

Autofluorescence in grey (488 channel) and staining in red (647 channel)

**A)** Antibody 40143-MM05 on non-infected Syrian hamster lungs shows aspecific interaction towards the outer regions of the lung lobe

**B)** Intense staining with antibody 40143-R001 in a Syrian hamster lung lobe, severe aspecific binding is observed

**C)** Aspecific interaction of antibody MA1-7401 in a lung lobe whereby similar bronchiolar structures as infected hamster lungs are stained

### Supplementary Fig 5

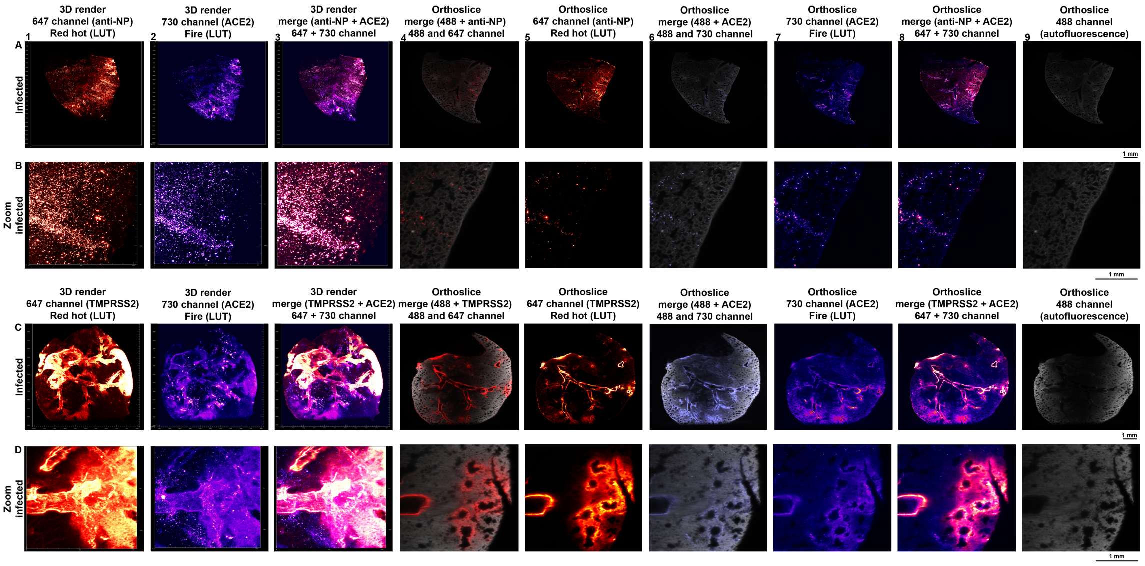


**Supplementary Fig 5. Supplementary analyses of NP, ACE2 and TMPRSS2 in a 4dpi Syrian hamster lung, related to figure 5.**

**A)** Top-to-bottom gradient of anti-NP stain, signal in the secondary bronchi extending towards the tertiary bronchi. A bottom-to-top gradient for ACE2 stain, with signal in the secondary and various tertiary bronchi. Overlap of anti-NP and anti-ACE2 in tertiary bronchi and bronchioles. 0.63 zoom, voxel resolution X,Y,Z: 4.79 µm, 4.79 µm, 5 µm.

**B)** anti-NP and ACE2 foci overlayed, severe overlap is visualized in the alveoli. 6.3 zoom, voxel resolution X,Y,Z: 0.48 µm, 0.48 µm, 2 µm.

**C)** TMPRSS2 expression profile, predominant staining in the primary, secondary and tertiary bronchi with signal spreading towards the bronchioles. ACE2 and TMPRSS2 superimpose in the primarily in the tertiary bronchi and bronchioles 0.63 zoom, voxel resolution X,Y,Z: 4.79 µm, 4.79 µm, 5 µm.

**D)** Overlap of TMPRSS2 and ACE2 only in the bronchioles, whilst ACE2 expression is also present in the alveoli. 6.3 zoom, voxel resolution X,Y,Z: 0.48 µm, 0.48 µm, 2 µm**.**

**Row 1)** 3D render 647 channel (Red hot false color).

**Row 2)** 3D render 730 channel (Fire false color).

**Row 3)** merge 3D render 647 channel (Red hot false color) and 730 channel (Fire false color).

**Row 4)** orthoslice merge 488 and 647 channel.

**Row 5)** Orthoslice 647 channel (Red hot false color).

**Row 6)** orthoslice merge 488 and 730 channel.

**Row 7)** orthoslice 730 channel (Fire false color).

**Row 8)** merge orthoslice 647 channel (Red hot false color) and 730 channel (Fire false color).

**Row 9)** orthoslice 488 channel (autofluorescence)

#### Supplementary Fig 6

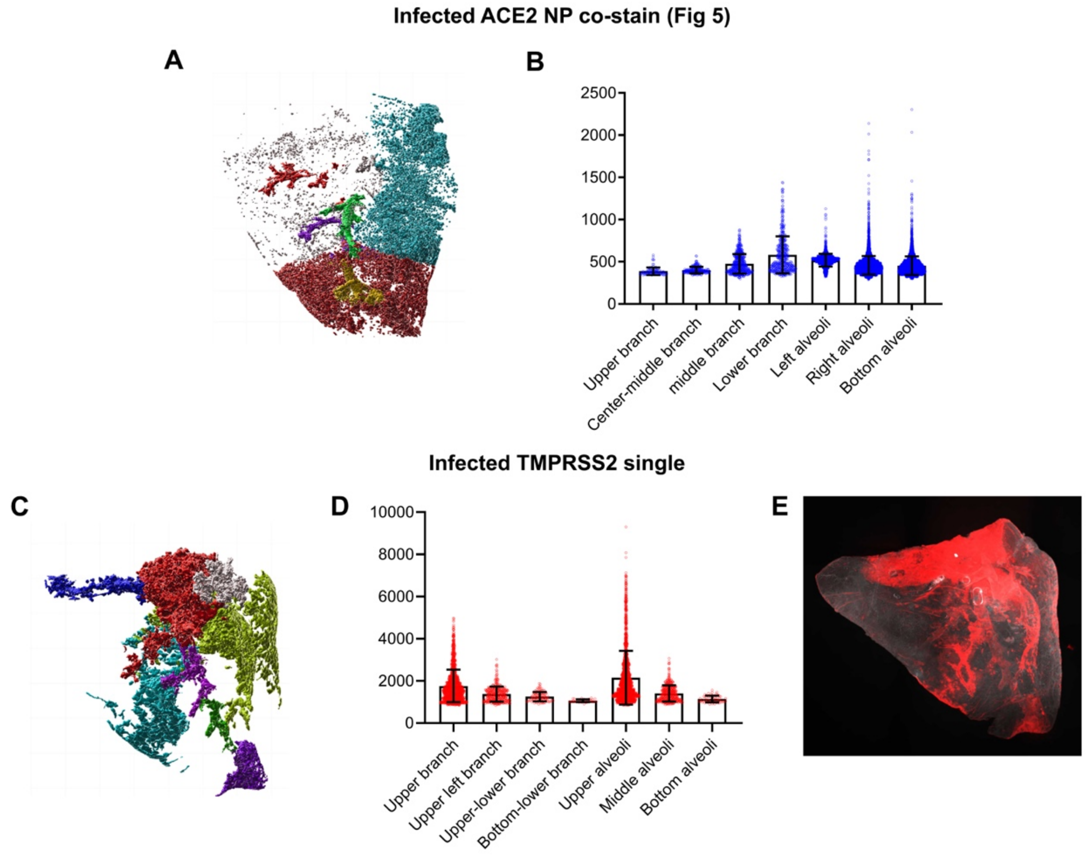


**Supplementary Fig 6. Supplementary ACE2 and TMPRSS2 signal quantifications in Syrian hamster lung lobes.**

**A)** Positions chosen for quantification in an infected Syrian hamster lung lobe from figure 5 A, different branches and alveoli are indicated.

**B)** Quantification of ACE2 signal in an infected Syrian hamster lung lobe from Fig 5A.

**C)** Positions chosen for quantification in an infected Syrain hamster lung stained with antibodies to TMPRSS2 (sample v20-441), different branches and alveoli are indicated.

**D)** Quantification of TMPRSS2 signal in an infected Syrian hamster lung lobe.

**E)** Significant TMPRSS2 expression in primary, secondary and tertiary bronchi with minor alveolar presence.
